# Simultaneous cortical, subcortical, and brainstem mapping of sensory activation

**DOI:** 10.1101/2024.04.11.589099

**Authors:** Neha A. Reddy, Rebecca G. Clements, Jonathan C. W. Brooks, Molly G. Bright

**Author notes:** **Corresponding Author:** Neha A. Reddy 645 N. Michigan Ave., Suite 1100 Chicago, IL 60611 USA.

## Abstract

Non-painful tactile sensory stimuli are processed in the cortex, subcortex, and brainstem. Recent functional magnetic resonance imaging (fMRI) studies have highlighted the value of whole-brain, systems-level investigation for examining pain processing. However, whole-brain fMRI studies are uncommon, in part due to challenges with signal to noise when studying the brainstem. Furthermore, the differentiation of small sensory brainstem structures such as the cuneate and gracile nuclei necessitates high resolution imaging. To address this gap in systems-level sensory investigation, we employed a whole-brain, multi-echo fMRI acquisition at 3T with multi-echo independent component analysis (ME-ICA) denoising and brainstem-specific modeling to enable detection of activation across the entire sensory system. In healthy participants, we examined patterns of activity in response to non-painful brushing of the right hand, left hand, and right foot, and found the expected lateralization, with distinct cortical and subcortical responses for upper and lower limb stimulation. At the brainstem level, we were able to differentiate the small, adjacent cuneate and gracile nuclei, corresponding to hand and foot stimulation respectively. Our findings demonstrate that simultaneous cortical, subcortical, and brainstem mapping at 3T could be a key tool to understand the sensory system in both healthy individuals and clinical cohorts with sensory deficits.

## Introduction

Processing of non-painful tactile sensory stimuli involves brain regions across the cortex, subcortex, and brainstem. The dorsal column pathway conveys sensation of light touch, vibration, brush and proprioception; it ascends ipsilaterally in the spinal cord, forming synaptic connections in the brainstem medulla before decussating to project to the thalamus, and finally terminating in sensorimotor cortex. The spinocerebellar tracts, parallel pathways also involved in proprioception, carry stretch-related signals from muscles via the spinal cord to the cerebellum. Tactile sensory processing throughout the sensory network is affected in a wide variety of conditions, including stroke, diabetes, autism spectrum disorders, Parkinson’s disease and by normal aging (Carey et al. 2011; Brodoehl et al. 2013; He et al. 2021; Lorenzini et al. 2021; Chitneni et al. 2022; Croosu et al. 2023; Ma et al. 2023). The changes in tactile detection and discrimination experienced in these conditions can affect mobility, self-care, and in the case of autism spectrum disorders, make it difficult to engage in social activities like childhood play (Welmer et al. 2007; He et al. 2021). A systems view of non-painful sensory processing is therefore a critical tool to study these disorders in both basic and translational research.

This full systems study requires investigation of brain function at the cortical surface and in deeper structures, which is possible through blood oxygenation level dependent (BOLD) functional magnetic resonance imaging (fMRI). Increasingly, fMRI studies have been highlighting the importance of cortical and subcortical/brainstem interactions in understanding brain systems. Recent functional connectivity studies including subcortical and brainstem regions have found that these extra-cortical areas are integrated with cortical function (Hansen et al. 2023; Hirsch and Wohlschlaeger 2023). In particular, cortical-brainstem fMRI has proved to be beneficial in the studies of pain and auditory processing, two systems that involve key brainstem nuclei. Pain studies utilizing simultaneous cortical-brainstem fMRI have shown that brainstem circuits are involved in modulation of pain pathways (Oliva et al. 2021; Oliva et al. 2022). Whole brain studies of auditory processing have also furthered our understanding of auditory subcortical and brainstem nuclei and their temporal correlations with cortical regions (Griffiths et al. 2001; Sigalovsky and Melcher 2006; Schönwiesner et al. 2007). In addition to these studies of pain and auditory processing in healthy individuals, the opportunities for discovery in clinical populations, including chronic pain and Parkinson’s disease, with brainstem fMRI have been detailed previously (Tracey and Iannetti 2006; Sclocco et al. 2018).

Despite the potential utility of systems-level investigation, fMRI studies of non-painful tactile sensory activation in humans have primarily focused on cortical areas, with a few studies studying only specific subcortical regions in isolation. In the cortex, the primary and secondary somatosensory areas are activated, and location of the stimulus has been shown to correspond with the somatosensory homunculus (Disbrow et al. 2000; Ruben et al. 2001; Del Gratta et al. 2002; Eickhoff et al. 2008; Gordon et al. 2023). The thalamus and cerebellum have also been independently studied. In the thalamus, the ventral posterolateral nucleus has been linked to processing of tactile stimuli (Gilman 2002; Charyasz et al. 2023; Habig et al. 2023). In the cerebellum, upper and lower extremity movements have been observed to activate lobules V/VI/VIIIa/VIIIb and I-IV/V/VIIIb/XI, respectively (Grodd et al. 2001; Ashida et al. 2019).

In contrast to studies of cortical and subcortical brain regions, fMRI studies of brainstem activity are much more limited, and the ability of brainstem fMRI to identify specific non-painful tactile sensory nuclei has not been demonstrated. In the tactile sensory system, the key brainstem regions are the cuneate and gracile nuclei of the medulla. The bilateral cuneate nuclei are involved in upper extremity and trunk sensory processing, and the more medial gracile nuclei are involved in lower extremity sensory processing (Vanderah and Gould 2015). The ability to localize and differentiate these nuclei with fMRI would allow us to better characterize regional sensory function in healthy and clinical populations at the brainstem level. fMRI studies using signal intensity enhancement by extravascular protons (SEEP) have applied brushing and painful stimuli to identify activity in the general region of the cuneate and gracile nuclei (Ghazni et al. 2010; Cahill and Stroman 2011). While a few BOLD fMRI studies have detected activity consistent with the cuneate nucleus using a finger tapping stimulus (Pattinson, Governo, et al. 2009; Faull et al. 2015), no study, to our knowledge, has mapped activity specific to the gracile nucleus or identified and differentiated these small but critical adjacent nuclei using fMRI.

A major constraint is that brainstem fMRI is challenged by high physiological noise, small nuclei sizes, and relative distance from radiofrequency receive coils (Brooks et al. 2013; Beissner 2015). While a limited number of studies have probed subcortical activity during non-painful sensory stimuli, these challenges have so far precluded a full systems study of the cortical, subcortical, and brainstem regions involved in sensory processing. Existing techniques to address the challenges of brainstem fMRI include using a restricted field of view (DaSilva et al. 2002; Pattinson, Mitsis, et al. 2009; Kubina et al. 2010; Faull et al. 2015; Matt et al. 2019), moving from 3T to 7T MRI (Hahn et al. 2013; Faull et al. 2015; Priovoulos et al. 2018; Sclocco et al. 2018), and implementing physiological denoising strategies like RETROICOR (Glover et al. 2000; Limbrick-Oldfield et al. 2012; Brooks et al. 2013; Sclocco et al. 2020; Oliva et al. 2021). While restricted field-of-view studies can enable greater spatial resolution, they limit the scope of brain systems-level investigation. 7T MRI systems also enable greater sensitivity and spatial resolution, but are not as widely available as 3T MRI systems, limiting their utility in both research and clinical settings. In addition, the issues of physiological noise (Krüger and Glover 2001; Triantafyllou et al. 2005) and susceptibility-induced distortions (Gizewski et al. 2014) are increased at 7T.

Another denoising strategy that has been shown to improve data quality in subcortical and brainstem regions is multi-echo independent component analysis (ME-ICA). This technique involves acquiring data at multiple echo times; during ICA denoising, multi-echo information can help classify components by quantifying the likelihood that each component is related to the true BOLD signal (e.g., T2* effects). Thus, ME-ICA provides a data-driven approach to identifying components associated with physiological pulsations, field changes, and other non-BOLD artifacts, such as those related to participant movement. Previous studies have demonstrated a temporal signal-to-noise ratio (tSNR) increase in the brainstem with both multi-echo fMRI acquisition and ME-ICA (Kundu et al. 2012; Dipasquale et al. 2017; Maugeri et al. 2018; Turker et al. 2021; Beckers et al. 2023), though this strategy has not yet been used to identify task activation in the brainstem. An added benefit of ME-ICA compared to physiological denoising methods like RETROICOR is the ability to decrease effects of task-correlated confounds and improve activation estimates (Evans et al. 2015; Gonzalez-Castillo et al. 2016; Lombardo et al. 2016; Cohen et al. 2018; Cohen and Wang 2019; Cohen, Jagra, Visser, et al. 2021; Cohen, Jagra, Yang, et al. 2021; Cohen, Chang, et al. 2021; Moia et al. 2021; Reddy et al. 2024). Task-correlated confounds like motion can be increased in clinical populations (Seto et al. 2001; Reddy et al. 2024), making ME-ICA a particularly valuable tool in using subcortical and brainstem fMRI to studying sensory deficits.

To address the aforementioned challenges and enable full brain mapping of the non-painful tactile sensory system, we implemented a targeted scan protocol that provides a whole-brain field of view, sufficient in-plane spatial resolution in the brainstem, and ME-ICA denoising for improved data quality. In a healthy population, we test our ability to detect and localize activity at each level of the sensory system and probe our sensitivity and specificity to activation in the brainstem.

## Materials and Methods

### Participants

This study was approved by the Northwestern University Institutional Review Board, and all participants provided written, informed consent. Sixteen right-handed, healthy participants with no known history of neurological or vascular disorders were scanned on a Siemens 3T Prisma MRI system with a 32-channel head coil. One participant was excluded from analysis due to an incidental finding, so data from fifteen of the participants are included (6M, 26 ± 3 years). A structural T1-weighted multi-echo MPRAGE image was collected using parameters adapted from Tisdall and colleagues (2016): TR = 2.17 s, TEs = 1.69/3.55/5.41 ms, TI = 1.16 s, FA = 7°, FOV = 256 x 256 mm^2^, and voxel size = 1 x 1 x 1 mm^3^. For four participants, a 64-channel head coil was used to acquire the MPRAGE image (Supplementary Table 1). The three echo images were combined using root-mean-square. A gradient-echo field map was collected at the beginning of each scan session for use in distortion correction: TR = 0.645 s, TEs = 4.92/7.38 ms, FA = 50°, voxel size = 3 x 3 x 3 mm^3^, and phase encoding direction P >> A. Functional scans were collected using a multiband multi-echo gradient-echo echo planar imaging sequence provided by the Center for Magnetic Resonance Research (CMRR, Minnesota): TR = 2.2 s, TEs = 13.4/39.5/65.6 ms, FA = 90°, MB factor = 2, GRAPPA = 2, voxel size = 1.731 x 1.731 x 4 mm^3^, 44 slices, phase encoding direction A >> P, field of view 180 mm, matrix size 104 x 104, and 235 volumes (Moeller et al. 2010; Setsompop et al. 2012). Axial slices were aligned perpendicular to the base of the fourth ventricle to maximize both in-plane resolution in the brainstem and whole-brain coverage.

### Sensory stimuli

During each functional scan, a non-painful sensory stimulus was applied to the participants’ hand or foot. An investigator (N.A.R.) manually brushed the hand or foot using a brush with stiff bristles, applied at a rate of 1 Hz for 12 repeats of 20-s brush / 20-s rest. Task and brush-rate timings were visually cued to the investigator on a screen. To maximize the number of sensory fibers activated (Corniani and Saal 2020), the brush was applied to the glabrous skin of the palm and fingers during the hand-stimulus scans and the lateral sole and toes during the foot-stimulus scans, across a distance of ∼130-160 mm; an overview of the sensory pathways involved in this stimulation is shown in Figure 1. Ten participants (4M, 26 ± 3y) underwent two functional scans each during stimulation of the right hand and left hand. Ten participants (5M, 25 ± 3y) underwent two functional scans during stimulation of the right foot. A detailed list of functional scans acquired for each participant is provided in Supplementary Figure 1.

**Figure 1.**
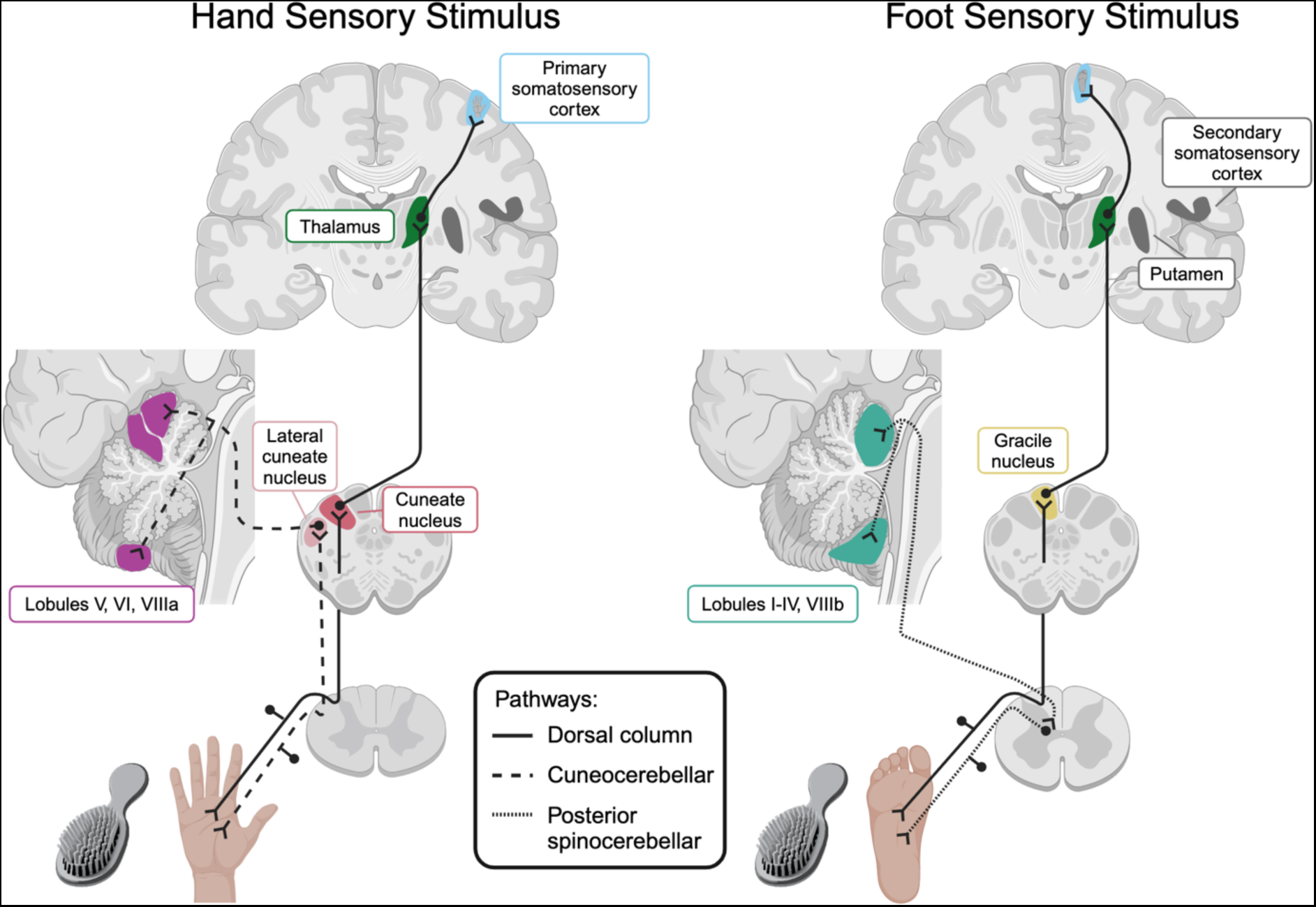
Pathways involved in hand and foot non-painful sensory stimulation: dorsal column and spinocerebellar. The dorsal column pathway synapses in the medulla, the thalamus, and the primary somatosensory cortex. In the medulla, the hand dorsal column pathway synapses in the cuneate nucleus, and the foot dorsal column pathway synapses in the gracile nucleus. The spinocerebellar tracts involved in tactile sensation and proprioception are the cuneocerebellar (upper extremity) and posterior spinocerebellar (lower extremity) pathways. The cuneocerebellar pathway synapses in the lateral cuneate nucleus of the medulla and enters the cerebellum via the inferior cerebellar peduncle. The posterior spinocerebellar pathway synapses in Clarke’s nucleus in the spinal cord before entering the cerebellum via the inferior cerebellar peduncle. Additional regions that have also been implicated in sensory processing are highlighted in dark gray. Created with BioRender.com.

### Structural MRI pre-processing

T1-weighted images for each subject were processed with FSL’s (Jenkinson et al. 2012) fsl_anat, which performs bias field correction and brain extraction.

### Functional MRI pre-processing

FSL (version 6.0.7.2) (Jenkinson et al. 2012) and AFNI (version 23.2.12) (Cox J.S. 1996) tools were used for fMRI preprocessing. The first 10 volumes of each echo timeseries were removed to allow for steady-state magnetization to be attained. Head-motion realignment was estimated for the first echo data, with reference to the Single Band reference image taken at the start of the scan (3dVolreg, AFNI), and then applied to all echo timeseries (3dAllineate, AFNI). All images were brain extracted (bet, FSL) and distortion corrected (FUGUE, FSL). Tedana (version 23.0.1) (Dupre et al. 2021; Ahmed et al. 2023) was used to calculate a T_2_*-weighted combination of the three echo datasets, producing the optimally combined (ME-OC) fMRI dataset. Multi-echo independent component analysis (ME-ICA) was performed on the ME-OC fMRI data using tedana. The resulting components were manually classified (accepted as signal of interest or rejected as noise), using the criteria described in (Reddy et al. 2024) and aided by Rica (Uruñuela 2021). The ME-OC timeseries at each voxel X was converted to signal percentage change for further analysis. No smoothing was applied.

### Subject-level model

A sensory task regressor was created by convolving the timing of the sensory stimulus with the canonical double-gamma hemodynamic response function. For each subject and stimulus, the ME-OC signal from each voxel was processed using a general linear model that incorporated both functional scan sessions and included six motion parameters from volume realignment, up to fourth-order Legendre polynomials, the sensory task regressor, and the rejected ME-ICA components (AFNI, 3dREMLfit).

### Whole-brain sensory activation group analysis

Group-level activation maps were calculated for each sensory stimulus (right hand, left hand, and right foot). Beta coefficient and t-statistic maps for the sensory stimulus regressors were converted to MNI space by applying a concatenated spatial transformation of the functional images to the subject’s T1-w structural image (epi_reg with field map unwarping, FSL) and then from this image to the 1-mm MNI template (FLIRT and FNIRT, FSL). The functional to anatomical and anatomical to standard registrations were visually inspected to ensure proper alignment of the lateral ventricles and gray matter-white matter borders. Group-level analysis across the whole brain was performed using AFNI’s 3dMEMA (Chen et al. 2012), using right-sided one-sample t-tests for each sensory stimulus. Group-level maps were thresholded at p < 0.001 and clustered at α < 0.05 (3dFWHMx, 3dClustSim, 3dClusterize, AFNI). The right-foot group-level map was also thresholded at p < 0.005 (without clustering) to demonstrate activation in key thalamus regions that were below the clustering threshold.

### Brainstem-specific sensory activation group analysis

Brainstem-specific group analysis was performed to increase sensitivity and specificity to activation in the brainstem, as done previously (Brooks et al. 2017; Oliva et al. 2021; Oliva et al. 2022). Threshold-free cluster enhancement (Smith and Nichols 2009) was performed within a mask of the lower medulla that contained the cuneate and gracile nuclei targeted in this study. The medulla mask was created by thresholding the brainstem region from the Harvard-Oxford Subcortical Atlas (Frazier et al. 2005) at 50%, then manually removing axial slices superior to the cuneate/gracile nuclei and extending the axial slices to the inferior limit of the MNI template. Analysis was performed with FSL’s non-parametric permutation test RANDOMISE (Nichols and Holmes 2001) and right-sided one-sample t-tests for each sensory stimulus. The maximum number of permutations using ten inputs was used (1024). Group-level maps are reported using family-wise error correction (TFCE_corrp) and thresholded at p < 0.05.

## Results

For each sensory stimulus, all subjects successfully completed the task paradigm during two fMRI scans. Significant group-level activation was mapped throughout anticipated regions of the cortex, subcortex, and brainstem.

### Cortical activation

At the cortical level, the dorsal column pathway synapses in the primary somatosensory cortex (S1), located in the postcentral gyrus (Figure 1). In our dataset, significant group-level cortical activation was found in the contralateral S1 for all sensory stimuli (Figure 2). Right-foot activity was located next to the inter-hemispheric fissure, with right-hand activity found laterally, aligning with the respective regions of the sensory homunculus (Figure 3). Secondary somatosensory cortex (S2) activity was also seen for all stimuli; hand stimuli demonstrated bilateral activity and the foot stimulus demonstrated only contralateral activity. Foot activation was observed medial to hand activation in S2 (Figure 3). For the right-hand task, a small region of activation was also seen next to the inter-hemispheric fissure (Supplementary Figure 1).

**Figure 2.**
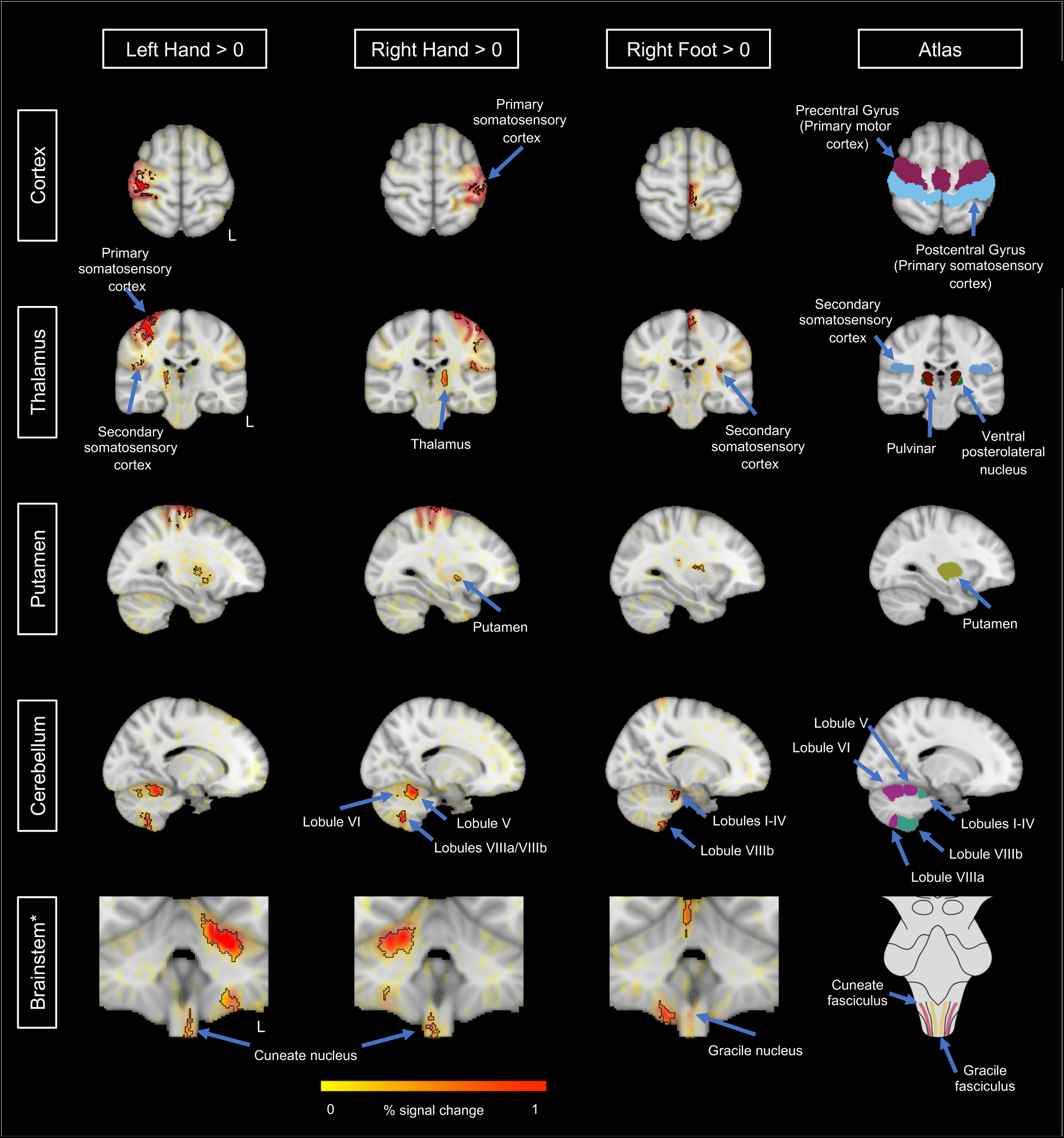
Group results for whole-brain analysis with 3dMEMA, highlighting activity in the cortex, thalamus, putamen, cerebellum, and brainstem. Opacity of beta coefficients is modulated by the t-statistic, as recommended by Taylor and colleagues (2023). Significant clusters are outlined in black, found by thresholding at p < 0.001 and clustering at α < 0.05. Left-hand, right-hand, and right-foot stimuli demonstrated significant positive clusters in primary/secondary somatosensory cortices, primary motor cortex, thalamus, putamen, and cerebellum. Atlas regions are also shown for comparison: pre- and post-central gyrus from the Harvard-Oxford Cortical Atlas (Desikan et al. 2006), thresholded at 25% to highlight the border between the gyri; secondary somatosensory cortex from the Julich Histological Atlas (Amunts et al. 2020), thresholded at 50%; pulvinar and ventral posterolateral nuclei of the Saranathan thalamic atlas (Saranathan et al. 2021); putamen from the Harvard-Oxford Subcortical Atlas (Frazier et al. 2005), thresholded at 50%; cerebellum lobules I-IV, V, VI, and VIIIa/b from the SUIT probabilistic cerebellum atlas (Diedrichsen et al. 2009; Diedrichsen et al. 2011); and brainstem region original line drawing adapted from Duvernoy (1995). *Brainstem-specific analysis is shown in Figure 4. Additional views for the right-hand stimulus are shown in Supplementary Figure 1.

**Figure 3.**
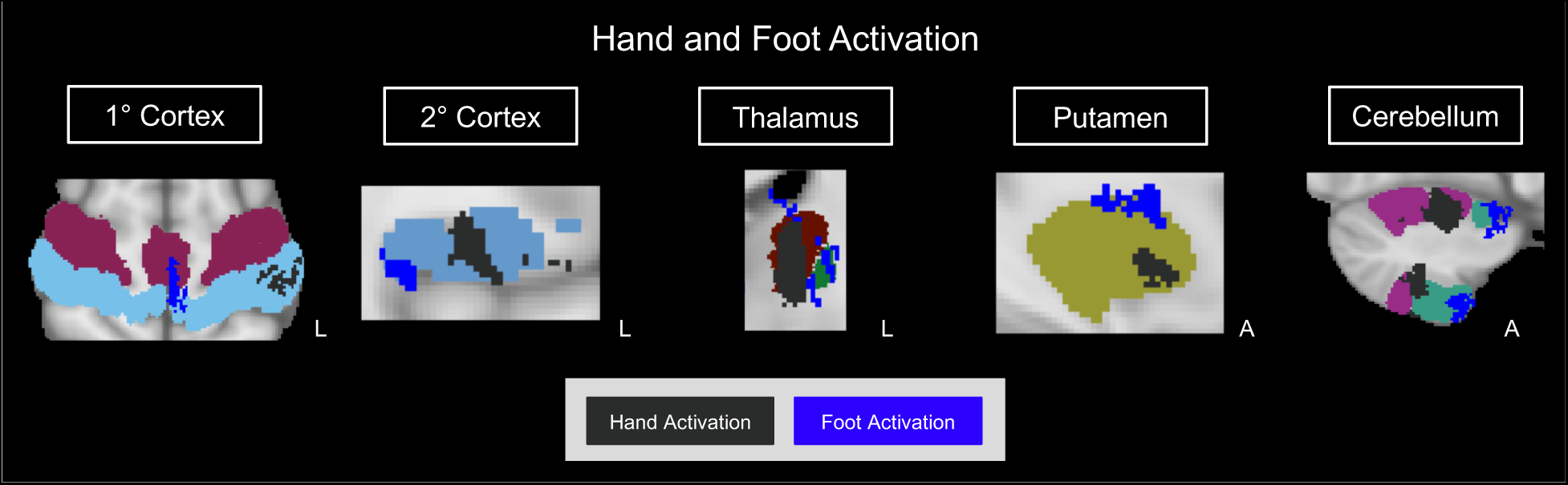
Group results for whole-brain analysis with 3dMEMA, comparing right-hand and right-foot activation. Significant clusters of activation for the right hand are shown in black and for the right foot shown in blue. For the thalamus only, right-foot activation was visualized with threshold p < 0.005 to demonstrate regions that were below the clustering threshold used for all other brain activity. Relevant atlas regions shown in Figure 2 are displayed for context.

In addition to the expected regions of sensory cortex activation, all stimuli had significant activation in the contralateral primary motor cortex (M1) in the precentral gyrus (Figure 2, Supplementary Figure 1). Similar to somatosensory cortex activity, locations of M1 activity aligned with regions of the motor homunculus, just anterior to S1 activity.

### Subcortical activation

At the subcortical level, the dorsal column pathway synapses in the thalamus (Figure 1). Both hand stimuli demonstrated significant activation in the contralateral thalamus, in a region roughly aligning with the pulvinar nucleus. While it did not pass the clustering threshold, the foot stimulus also showed activity in the contralateral thalamus, medial to the hand stimuli and aligning with the ventral posterolateral nucleus (Figure 3).

The spinocerebellar pathway is another component of the sensory system that synapses in the cerebellum after ascending the spinal cord. Both hand stimuli demonstrated activity in the ipsilateral cerebellum, in lobules V/VI and VIIIa/b. The foot stimulus demonstrated ipsilateral activity in lobules I-IV and VIIIb (Figure 2). The right-hand stimulus additional demonstrated activity in contralateral lobules VIIIa/b.

While not associated with the dorsal column or spinocerebellar pathways, activity was also detected in the contralateral putamen for all stimuli. Foot putamen activity was located superior to hand putamen activity (Figure 3).

### Brainstem activation

In the brainstem, the dorsal column pathway synapses in the cuneate and gracile nuclei of the medulla. Brainstem activation was investigated with whole-brain and brainstem-specific group analyses to increase sensitivity to activation in the region (Brooks et al. 2017; Oliva et al. 2021; Oliva et al. 2022). In the whole-brain group analysis, brainstem activation was detected in the ipsilateral medulla for right- and left-hand stimuli (Figure 2). However, areas of significant activation encompassed a large region, not specific to the cuneate nuclei associated with hand sensation. At the whole-brain level, no significant cluster of activation related to stimulation of the right-foot was detected in the brainstem.

Brainstem-specific group analysis was performed to increase our sensitivity and specificity to activation in this region. Based on previous knowledge of the brainstem nuclei involved in sensory stimuli, this group analysis was performed within a mask of the lower medulla that included all axial slices expected to contain the right and left cuneate and gracile nuclei. Significant clusters of activation related to the right-hand, left-hand, and right-foot stimuli were detected in the medulla (Figure 4). For all stimuli, significant activation was found ipsilateral to the stimulus, as expected based on the dorsal column pathway. There was one voxel of overlap between significant activation clusters found for right-hand and right-foot stimuli. Otherwise, significant activation clusters were distinct between stimuli. Right-hand activation in the cuneate nucleus was lateral and inferior to right-foot activation in the gracile nucleus, aligning with brainstem atlases of these nuclei (Paxinos et al. 2012; Adil et al. 2021).

**Figure 4.**
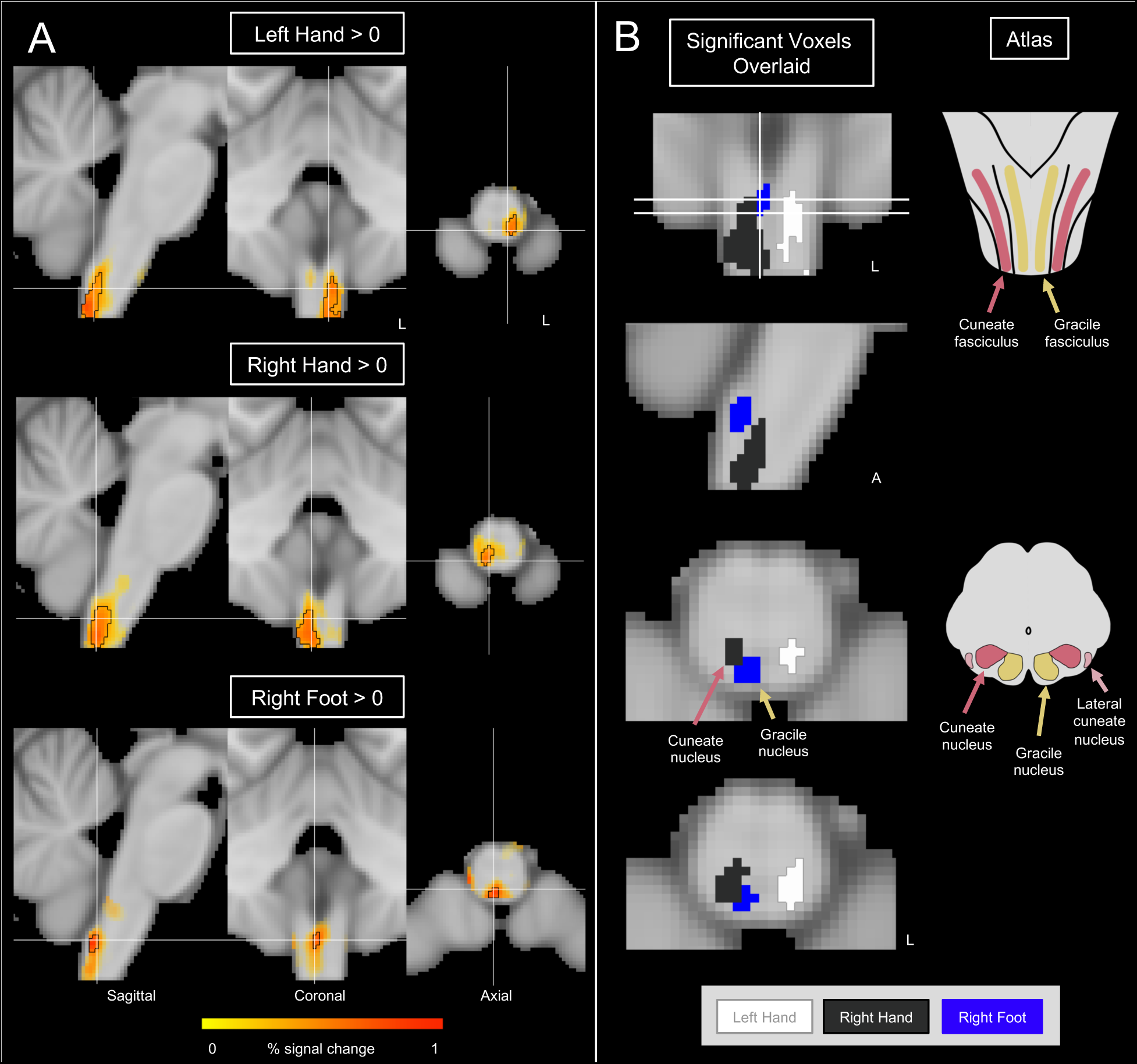
Group results for brainstem-specific analysis with FSL’s RANDOMISE. (A) Left-hand, right-hand, and right-foot stimuli activation shown separately. Opacity of beta coefficients is modulated by the TFCE- and FWE-corrected p-value. Significant voxels (p < 0.05) are outlined. (B) Significant voxels (p < 0.05) for left-hand, right-hand, and right-foot stimuli overlaid; there is no overlap of significant voxels for each stimulus in the depicted slices. Left- and right-hand stimuli demonstrated significant positive clusters in the ipsilateral cuneate nuclei. The right-foot stimulus demonstrated a significant positive cluster in the ipsilateral gracile nucleus. Original line drawings of brainstem atlas regions adapted from Duvernoy (1995).

## Discussion

In this study, we mapped the tactile sensory system across the cortex, subcortex, and brainstem, using multi-echo fMRI at 3T and a whole-brain field of view. We applied sensory stimulation to the right hand, left hand, and right foot in a cohort of healthy adults to assess our ability to identify and differentiate activity related to each stimulus. We performed both whole-brain and brainstem-specific analyses to take advantage of our large field-of-view while also enhancing sensitivity to activity in the brainstem, which exhibits lower signal to noise characteristics. In sensory regions across the cortex, subcortex, and brainstem, we were able to determine both appropriately lateralized activity for right- and left-hand stimuli and distinct areas of activity for right hand and foot stimuli. To our knowledge, this is the first time that task-fMRI techniques have been successfully used to discriminate the adjacent cuneate and gracile nuclei.

### Whole-brain sensory activation

Using our whole-brain analysis approach, we were able to identify specific regions of activation across the cortex and subcortex for all stimuli. We expected that regions of activation would align with the dorsal column and spinocerebellar pathways, which are involved in processing of non-painful tactile stimuli generated by our brushing protocol. To account for non-specific effects of sensory stimulation, such as attention, we additionally performed paired t-tests for the right-versus left-hand stimuli, which yielded similar results to those described below (Supplementary Figure 2).

The dorsal column pathway begins with stimulation of cutaneous sensory receptors; in our study, brushing of the palms of the hand and soles of the feet will have activated receptors in the glabrous skin, including Merkel endings, Meissner corpuscles, and Ruffini endings (Vanderah and Gould 2015). This pathway then ascends to synapse in the medulla. After decussating at the level of the medulla, the dorsal column synapses in the contralateral ventral posterolateral nucleus (VPL) of the thalamus, and finally terminates in the primary somatosensory cortex (Figure 1). We were able to identify and differentiate right and left, and hand and foot activity in each of these critical regions.

In the primary somatosensory cortex (S1), we observed activity that was contralateral and aligned with classical and modern depictions of the sensorimotor homunculus (Penfield and Rasmussen 1950; Gordon et al. 2023), with foot activity medial to hand activity (Figure 2). For the right-hand stimulus, a small region of activation was found in a region that aligns with the leg/foot region of the homunculus (Supplementary Figure 1); this may be due to passive movements of the right side of the body caused by brushing of the hand. Activity was also identified in the secondary somatosensory cortex (S2) (Figure 2). While not a part of the dorsal column pathway, the S2 receives sensory inputs from the S1 and the thalamus (Vanderah and Gould 2015). The right- and left-hand stimuli both resulted in bilateral S2 activity, similar to what has been described previously. For example, in a meta-analysis of tactile stimulus studies, Lamp and colleagues (2019) found that bilateral S2 activation was commonly seen when a tactile stimulus was applied to the right or left hand. The S2 has also been observed to have somatotopy, similar to the S1. Del Gratta and colleagues (2002) found that hand sensory activation was posterior and lateral to foot sensory activation in the S2. Aligning with these findings, we also observed that hand activity was lateral to foot activity in the contralateral S2 (Figure 3).

In the thalamus, we expected to detect activity in the contralateral VPL. While right-foot activity aligned with the VPL when compared to the Saranathan thalamic atlas (Saranathan et al. 2021), right-hand and left-hand activity were medial to the VPL, aligning more closely with the pulvinar region (Figure 2, Figure 3). Though not expected based on our knowledge of the dorsal column pathway, this finding does align with previous studies that observed pulvinar involvement during non-painful sensory processing (Golaszewski et al. 2006; Charyasz et al. 2023; Habig et al. 2023). In a study of hand and foot motor activity, Errante and colleagues (2023) also saw that hand activity was medial to foot activity and had more pulvinar involvement, consistent with our results. The pulvinar has been noted to be involved in multisensory integration, and thus may have been active during our tactile sensation stimulus (Froesel et al. 2021). The discrepancy between VPL and pulvinar activity in previous findings may also be influenced by potential misalignment of group-level results. Registration is a particularly important step when investigating subcortical regions with smaller nuclei, such as the thalamus. In this study, we used FSL’s FNIRT with visual inspection of registration results to ensure the quality of this processing step; other established and emerging registration methods have also been previously demonstrated (Avants et al. 2011; Lange et al. 2024). In addition, “precision mapping experiments” have shown that individual anatomy-function relationships may be unique (Gordon et al. 2017), and thus may provide a different lens to study thalamic nuclei activity compared to group-level analysis. We also note that our hand stimuli resulted in significant clusters in the thalamus, while foot activity in the region did not achieve significance at the same threshold, possibly due to the four-fold reduction in density of innervation in this region compared to the hand (Corniani and Saal 2020). We demonstrated the foot-related thalamic activity using other thresholding methods, but future work using a larger sample size or thalamus-specific analyses may improve sensitivity to this relatively smaller region of activation.

The spinocerebellar tracts ascend the spinal cord and pass through the inferior cerebellar peduncle to synapse in the cerebellum (Vanderah and Gould 2015). In our study, we found hand-related activity in ipsilateral lobules V, VI, and VIIIa/b and foot-related activity in ipsilateral lobules I-IV and VIIIb (Figure 2). Similar results have been demonstrated by several motor (Grodd et al. 2001; Spencer et al. 2007; Stoodley 2012; Ashida et al. 2019; Errante et al. 2023; Reddy et al. 2024) and sensory (Bushara et al. 2001; Takanashi et al. 2003; Ashida et al. 2019) studies of the cerebellum. Of these studies, only two use a whole-brain field of view and report additional results in regions outside the cerebellum, and both reflect motor task designs that may additionally involve sensory feedback (Errante et al. 2023; Reddy et al. 2024). Our finding of contralateral activation (during the right-hand stimulus only) has also been previously reported, and may be subject-specific (Bushara et al. 2001; Takanashi et al. 2003); again, future precision mapping experiments may further elucidate the significance of contralateral cerebellum usage in specific individuals.

In addition to activation of the expected sensory regions described above, we also observed clusters in the primary motor cortex (M1) and putamen. In the M1, activity for all stimuli was found in the contralateral precentral gyrus, directly anterior to activity in S1, aligning with the motor homunculus (Gordon et al. 2023) (Figure 2). Brushing of the hands and feet likely caused small passive or active movements that resulted in the observed M1 activity. With regards to putamen involvement, while the putamen is primarily considered to be related to motor activity, previous work has also implicated it in sensory processing (Goble et al. 2012; Vicente et al. 2012; Eckstein et al. 2020). In our study, we observed contralateral putamen activity related to all stimuli (Figure 2). We additionally found that foot-related activity was superior to hand-related activity in the putamen (Figure 3). Gerardin and colleagues (2003) demonstrated the same somatotopy in the putamen using motor-task fMRI with a restricted field of view.

### Sensitivity and specificity to activation in the brainstem

In the brainstem, we expected to observe activity in the ipsilateral cuneate and gracile nuclei of the medulla, related to hand and foot activity, respectively. Right-hand and left-hand stimulation may also activate the ipsilateral lateral cuneate nucleus, which is lateral to the cuneate nuclei (Figure 1), but we did not aim to discriminate the cuneate and lateral cuneate nuclei with our protocol. Our whole-brain analysis was able to detect significant clusters of ipsilateral activation in the medulla related to hand activity. While appropriate lateralization was detected, these clusters were more diffuse than expected. No significant clusters were found for the foot stimulus, although activity was similarly seen in the ipsilateral medulla. The reason for this discrepancy in significant detection of hand and foot activity may be due to the greater number and density of sensory fibers in the hand compared to the foot (Corniani and Saal 2020). Therefore, the same brushing stimulus may result in much more robust brain and brainstem activation when applied to the hand, compared to the foot.

In order to increase our sensitivity to brainstem activation across all stimuli in this study and assess our ability to discriminate the adjacent cuneate and gracile nuclei, we employed a brainstem-specific analysis within a mask of the lower medulla. This is similar to work by Brooks, Oliva, and colleagues, who used brainstem-specific analyses to characterize activity during painful thermal stimuli (Brooks et al. 2017; Oliva et al. 2021; Oliva et al. 2022). Using this method, we were able to identify distinct, significant regions of activation for all stimuli. The regions of activation for hand and foot stimuli aligned with existing brainstem atlases (Paxinos et al. 2012; Adil et al. 2021); foot activation in the gracile nuclei was medial and superior to hand activation in the cuneate nucleus. To our knowledge, no other study has differentiated these adjacent brainstem nuclei using fMRI. While a few studies have observed activity generally consistent with the cuneate nucleus (Pattinson, Governo, et al. 2009; Faull et al. 2015), no other study has explicitly assessed specificity of this activation or attempted to detect and distinguish activity in the gracile nucleus.

Of note, the cuneate and gracile nuclei of interest in our study are located in the inferior medulla, and the cuneate nuclei activation in our study reaches the most inferior slice of the standard MNI template. Capturing the full extent of cuneate nuclei activation may require extension of our analysis from the brainstem into the upper cervical spinal cord. While our structural and functional acquisitions extend into the upper cervical spinal cord, standard brain extraction tools, such as FSL’s bet used in this study, mask out regions inferior to the MNI template. Therefore, this extended analysis will require modified preprocessing steps, which may be possible through manual extension of automated brain masking functions, full analysis pipelines optimized for brainstem acquisitions (Oliva et al. 2022), and use of a combined brainstem/spinal cord standard atlas (De Leener et al. 2018). Although this analysis is outside the scope of the current study, such improvements will greatly benefit our characterization of brainstem function in neuroimaging data. In addition, similar to the whole-brain analysis, we performed brainstem-specific paired t-tests for the right-versus left-hand stimuli; these analyses yielded significant activation for the right-hand stimulus, but sub-threshold activation for the left-hand stimulus (Supplementary Figure 2). A greater sample size may enable reaching a target significance threshold for activation with a paired analysis.

### Limitations and considerations for future studies

Our study implemented a whole-brain field-of-view multi-echo fMRI acquisition at 3T to enable a simultaneous systems-level study of sensory processing. Out of the numerous sensory fMRI studies discussed previously, only a few use a whole-brain field-of-view to enable combined cortical-subcortical analysis, and none include brainstem analysis (Bushara et al. 2001; Golaszewski et al. 2006; Goble et al. 2012; Ashida et al. 2019). As mentioned previously, studies of painful (Oliva et al. 2021; Oliva et al. 2022) and auditory (Griffiths et al. 2001; Sigalovsky and Melcher 2006; Schönwiesner et al. 2007) stimuli have utilized a whole-brain approach, as well as one study of tongue motion (Corfield et al. 1999). However, these studies have not attempted to differentiate adjacent brainstem nuclei, as we have also successfully demonstrated in this study. In order to enable a whole-brain field-of-view with high resolution in the brainstem, we used a smaller in-plane resolution with oblique axial slices positioned perpendicular to the axis of the brainstem (1.731 x 1.731 mm) and a large z-axis resolution (4 mm) along the axis of the brainstem. Our acquisition protocol may not be feasible for studies that require higher z-axis resolution; a higher multiband factor may facilitate a smaller slice thickness, with the tradeoff of lower signal to noise and greater false-positive activations (Todd et al. 2016). Importantly, although our voxel dimensions are chosen to achieve in-plane specificity specifically in the brainstem, this anisotropic resolution did not limit our ability to detect anticipated cortical/subcortical activation.

In addition, we used multi-echo acquisition and denoising strategies to improve our data quality in subcortical and brainstem regions. In our study, both multi-echo acquisition and ME-ICA denoising provided significant and distinct improvements in temporal signal-to-noise ratio (tSNR) compared to a simulated single-echo approach (Supplementary Figure 1), in alignment with previous findings (Kundu et al. 2012; Dipasquale et al. 2017; Maugeri et al. 2018; Turker et al. 2021; Beckers et al. 2023). This tSNR increase, in addition to the reported decrease in between-subject variance with ME-ICA denoising, can reduce the required scan duration and sample size needed to detect activation (Murphy et al. 2007; Lombardo et al. 2016). A recent study by Mohamed and colleagues (2024) also demonstrated an increase in tSNR in specific brainstem nuclei using an optimized whole-brain acquisition protocol and physiological noise removal with the PhysIO toolbox (Kasper et al. 2017); our ME-ICA approach led to a median brainstem tSNR that is within the tSNR range of their tested nuclei. However, Mohamed and colleagues calculated their tSNR using nuclei in the midbrain and pons, while our tSNR measurement incorporated the entire brainstem, including the medulla. While a direct comparison is difficult due to this difference, these results demonstrate that the data quality afforded by our methods is comparable to other recent advancements in brainstem imaging.

ME-ICA denoising has also previously been shown to decrease effects of task-correlated confounds and improve reliability and stability of activation estimates (Evans et al. 2015; Gonzalez-Castillo et al. 2016; Lombardo et al. 2016; Cohen et al. 2018; Cohen and Wang 2019; Cohen, Jagra, Visser, et al. 2021; Cohen, Jagra, Yang, et al. 2021; Cohen, Chang, et al. 2021; Moia et al. 2021; Reddy et al. 2024). The target clinical populations for studying impaired sensorimotor function at the systems level are likely to have increased task-correlated artifacts (Seto et al. 2001; Reddy et al. 2024), and therefore would specifically benefit from a ME-ICA approach.

Our ME-ICA approach also has certain acquisition and analysis limitations. We chose an fMRI acquisition with three echo times to enable a whole-brain field of view and acceptable TR. However, acquiring more echo times may enable better optimal combination and calculation of ME-derived parameters used in ICA classification. In our data, the optimal echo time for the relevant regions ranged from ∼50-60 ms (Supplementary Figure 3). The highest echo time in our acquisition is 65.5 ms, which captures this window, but acquisition of additional higher echo times has the potential to enhance the utility of ME and ME-ICA techniques. A limitation in ME-ICA analysis is that the increase in subject-level model regressors decreases the degrees of freedom (Supplementary Figure 4). However, this decrease did not limit our ability to detect significant activation in the brainstem-specific group-level modeling. In addition, ME-ICA classification may require hands-on involvement; we found inspecting and manually revising the automatic component classification performed by tedana with study-specific criteria to be useful, as the default settings may not be appropriate for every study. Updates being currently developed in tedana will allow for creation of tailored ME-ICA classification pipelines that can mitigate manual classification time when extending the technique to larger sample sizes and clinical applications (Ahmed et al. 2023).

Although we did not directly incorporate physiological noise removal in this study, physiological denoising has previously been shown to improve data quality in brainstem fMRI (Harvey et al. 2008; Mohamed et al. 2024). ME-ICA is a more general approach that may indirectly correct for physiological noise, but the interactions between ME-ICA and physiological noise removal tools, such as RETROICOR (Glover et al. 2000) and the PhysIO toolbox (Kasper et al. 2017), have not been studied. While the unique explanatory characteristics of physiological noise correction compared to ICA have been demonstrated in specific instances (Krentz et al. 2023; Reddy et al. 2024), the effect may vary depending on the stimulus and cohort. Further work is necessary to determine the optimal usage of ME-ICA with existing physiological noise correction techniques.

Another key component of our fMRI acquisition was the 3T field strength. Higher field strengths have shown promise for brainstem fMRI due to increased spatial resolution and signal-to-noise ratio (Sclocco et al. 2018); however, the required MRI scanners are not yet widely available and may not be feasible for use by many research groups. As previously mentioned, physiological noise (Krüger and Glover 2001; Triantafyllou et al. 2005) and susceptibility-induced distortions around the brainstem (Gizewski et al. 2014) are also increased at 7T, potentially decreasing the utility of 7T systems for signal detection in the brainstem. Here, we demonstrate that a whole-brain analysis including sufficient brainstem resolution is possible at 3T, enabling wider systems-level study of the sensory system. The feasibility of 3T fMRI may be especially important for studies of clinical populations with sensory deficits, such as Parkinson’s disease and stroke. Understanding and delivering tactile sensation is also a critical component of developing brain-machine interface technologies that aim to restore motor control in amputees and individuals with spinal cord injury (Tabot et al. 2015; Collinger et al. 2018).

Clinical cohorts may be less able to travel to facilities with high-field scanners, and the possibility of 3T investigation enables broader involvement and collaboration of research groups for systems-level fMRI studies. 7T systems may also cause vertigo, nausea, and discomfort during scanning, making it difficult for clinical cohorts and children to tolerate scanning (Vargas et al. 2018). Higher specific absorption rates (SAR) caused by 7T imaging may increase constraints in populations with deep brain stimulation (DBS) implants (Larson et al. 2008; Vargas et al. 2018).

## Conclusions

In this study, we conducted the first simultaneous investigation of cortical, subcortical, and brainstem activity in response to tactile sensory stimulation and recorded using fMRI. Brainstem fMRI is particularly susceptible to poor signal to noise imaging data, which has historically been a significant challenge to performing such studies. We employed a targeted acquisition and multi-echo denoising strategy to enable a whole-brain field-of-view with high sensitivity to activation in the brainstem at 3T. We were able to identify specific areas of activation for non-painful hand and foot tactile stimuli in the cortex, thalamus, putamen, cerebellum, and medulla; and, for the first time, we demonstrated the ability of fMRI to differentiate adjacent sensory nuclei in the brainstem. Our findings demonstrate the feasibility of non-painful, sensory task-activation studies of the cortex, subcortex, and brainstem. Whole-brain/brainstem acquisitions permit concurrent sampling across the sensorimotor network, and offer the possibility to investigate disrupted sensory processing and connectivity in a wide range of clinical cohorts.

## CRediT author contribution statement

**Neha A. Reddy:** Conceptualization, Methodology, Software, Formal analysis, Investigation, Data curation, Writing – original draft, Writing – review & editing, Visualization, Project administration. **Rebecca G. Clements:** Investigation, Writing – review & editing. **Jonathan C. W. Brooks:** Conceptualization, Methodology, Writing – review & editing. **Molly G. Bright:** Conceptualization, Methodology, Writing – review & editing, Supervision, Project administration, Funding acquisition.

## Acknowledgements

This work was supported by the Center for Translational Imaging at Northwestern University and through the computational resources and staff contributions provided for the Quest high performance computing facility at Northwestern University, which is jointly supported by the Office of the Provost, the Office for Research, and Northwestern University Information Technology. Thank you to Julius P. A. Dewald, César Caballero-Gaudes, and Marta Bianciardi for their advice and guidance.

