## Supplementary Material for "Simultaneous cortical, subcortical, and brainstem mapping of sensory activation"

**Supplementary Table 1.** Scan information detailing whether a 32-channel or 64-channel head coil was used for the MPRAGE scan, and which functional scans were acquired in each participant. Ten participants completed the functional scans on two days; five participants completed the functional scans in one day. RH = Right Hand; LH = Left Hand; RF = Right Foot. *MPRAGE scan is missing one slice that does not contain any region of expected activation.

| Subject | MPRAGE | LH run 1 | LH run 2 | RH run 1 | RH run 2 | RF run 1 | RF run 2 |
|---|---|---|---|---|---|---|---|
| sub-01 | 32-ch | Day 1 | Day 2 | Day 1 | Day 2 | - | - |
| sub-02 | 32-ch | Day 1 | Day 2 | Day 1 | Day 2 | - | - |
| sub-03 | 32-ch | Day 1 | Day 2 | Day 1 | Day 2 | - | - |
| sub-04 | 32-ch | Day 1 | Day 2 | Day 1 | Day 2 | - | - |
| sub-05 | 32-ch | Day 1 | Day 2 | Day 1 | Day 2 | - | - |
| sub-06 | 32-ch | Day 1 | Day 2 | Day 1 | Day 2 | Day 1 | Day 2 |
| sub-07 | 32-ch | Day 1 | Day 2 | Day 1 | Day 2 | Day 1 | Day 2 |
| sub-08 | 32-ch | Day 1 | Day 2 | Day 1 | Day 2 | Day 1 | Day 2 |
| sub-09 | 32-ch | Day 1 | Day 2 | Day 1 | Day 2 | Day 1 | Day 2 |
| sub-10 | 32-ch | Day 1 | Day 2 | Day 1 | Day 2 | Day 1 | Day 2 |
| sub-11 | 64-ch | - | - | - | - | Day 1 | Day 1 |
| sub-12 | 64-ch | - | - | - | - | Day 1 | Day 1 |
| sub-13 | 64-ch | - | - | - | - | Day 1 | Day 1 |
| sub-14 | 64-ch | - | - | - | - | Day 1 | Day 1 |
| sub-15 | 32-ch* | - | - | - | - | Day 1 | Day 1 |

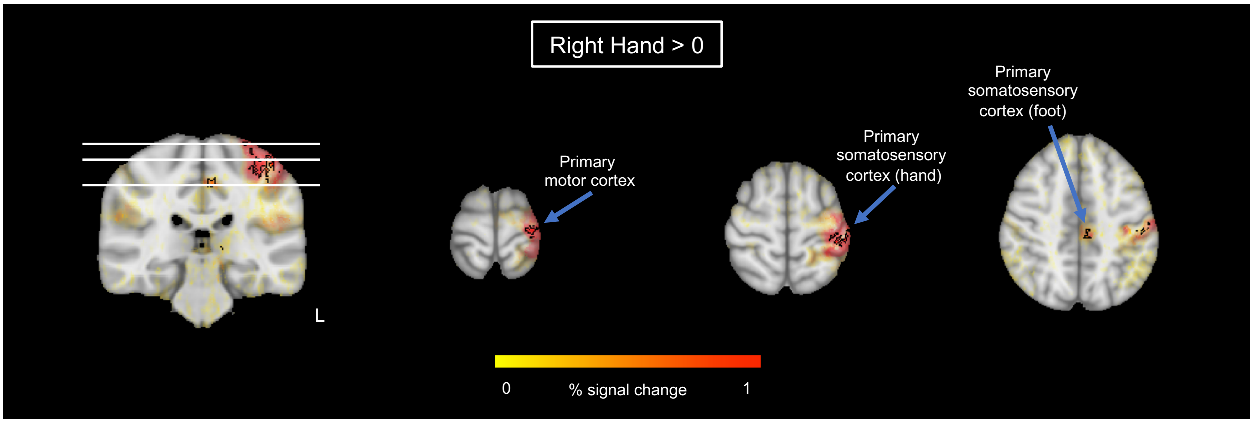


**Supplementary Figure 1.** Additional views of whole-brain group analysis for Right Hand > 0.

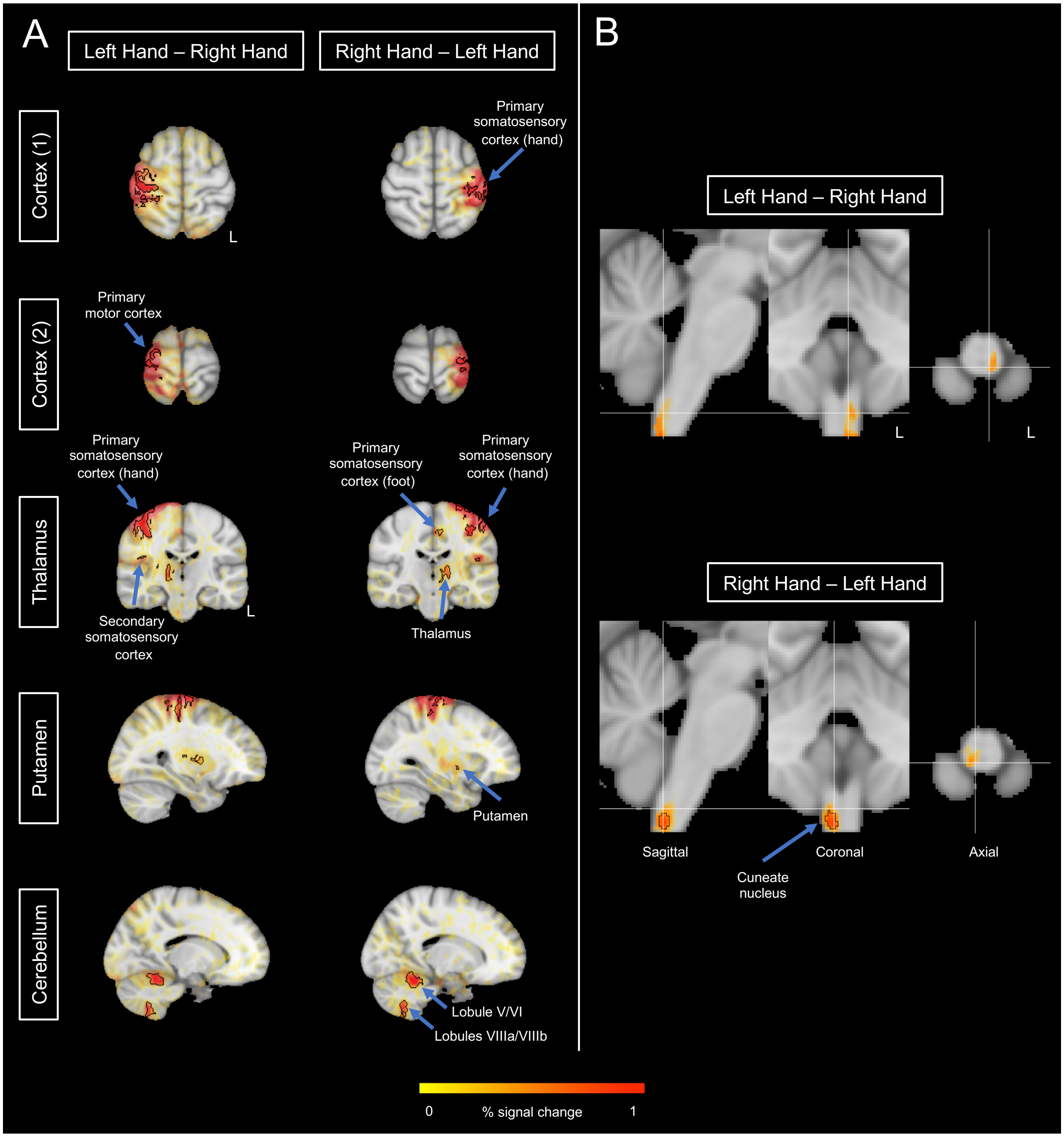


**Supplementary Figure 2.** Paired, right-sided t-tests for Left Hand – Right Hand and Right Hand – Left Hand. (A) Whole-brain group analysis using AFNI's 3dttest++ and cluster-based analysis. Opacity of beta coefficients is modulated by the t-statistic. Significant clusters are outlined in black, found by thresholding at $p < 0.001$ and clustering at $\boldsymbol{\alpha} < 0.05$. (B) Brainstem-specific group analysis using FSL RANDOMISE. Opacity of beta coefficients is modulated by the TFCE- and FWE-corrected p-value. Significant voxels ($p < 0.05$) are outlined. There were no significant voxels for the Left Hand – Right Hand condition.

**Temporal signal-to-noise ratio (tSNR) calculation**

The temporal signal-to-noise ratio (tSNR) of the ME-ICA denoised datasets was compared to the denoised ME-OC and single-echo (SE) datasets to understand how ME and ICA analyses affect data quality. SE and ME-OC analyses were performed for the purpose of tSNR comparison, and these data were not used for any further analyses in this paper.

The second echo data (TE=39.5s) were used as a surrogate for a standard SE fMRI acquisition. The first 10 volumes of each scan were removed to allow for steady-state magnetization. Head-motion realignment was computed with reference to the Single Band reference image taken at the start of the scan (3dVolreg, AFNI). After brain extraction (bet, FSL) and distortion correction (FUGUE, FSL), the SE timeseries were converted to signal percentage change for use in further analysis.

SE and ME-OC data were modeled with AFNI's 3dREMLfit, with models including six motion parameters from volume realignment, up to fourth-order Legendre polynomials, and the sensory task regressor. For the SE, ME-OC, and ME-ICA data, the residual timeseries from each model was used as the denoised dataset. The SE, ME-OC, and ME-ICA denoised datasets were reverted from signal percentage change for each voxel, in order to represent the true mean in tSNR analysis. Each voxel Y was converted as [Y*mean)]+mean(X), with X being the original voxel timeseries before signal percentage change conversion. tSNR of the denoised datasets was calculated for each voxel by dividing the mean of the timeseries by the standard deviation. A median tSNR was found within masks of the whole brain and of the brainstem. A brainstem mask was created for each scan using the Harvard-Oxford subcortical structural atlas brainstem map (Frazier et al. 2005) thresholded at 50%, manually extended to the inferior border of the MNI brain, then transformed to each scan's functional space. Significant differences in the tSNR between the SE, ME-OC, and ME-ICA models were tested as tSNR ~ Model + (1|Subject).

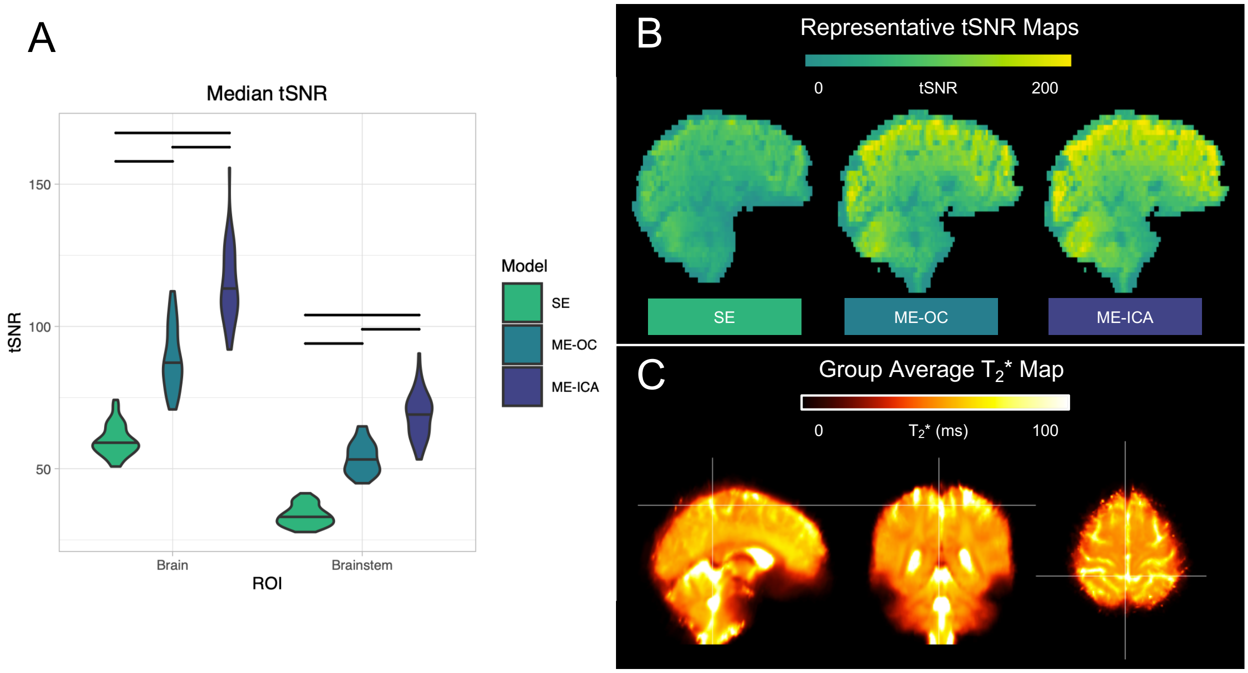


**Supplementary Figure 3.** (A) Median temporal signal-to-noise ratio (tSNR) in the whole brain and brainstem regions, compared between SE, ME-OC, and ME-ICA models. 60 datasets are included per model. Bars represent a significant difference in tSNR between models ($p < 0.0001$, Bonferroni-corrected). (B) Representative tSNR maps from one scan for each model, shown in functional space. (C) (A) $T_2^*$ map averaged across all subjects and scans. One subject-level $T_2^*$ map per subject was transformed to standard space, then averaged across all subjects.

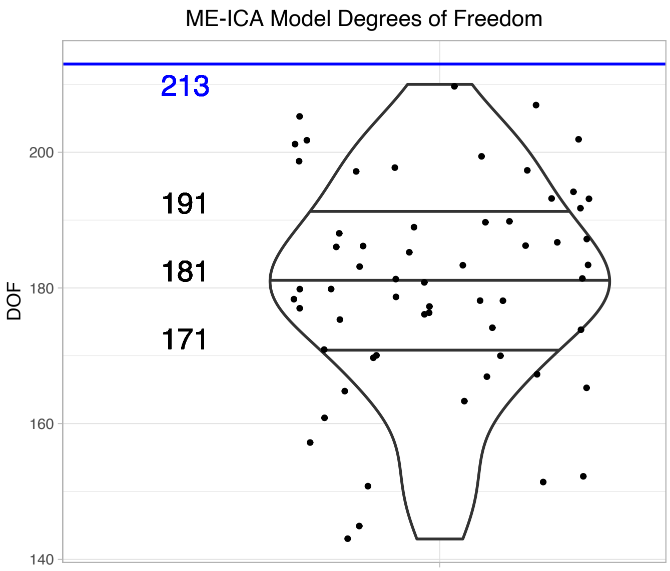


**Supplementary Figure 4.** Degrees of freedom (DOF) in ME-ICA models. All SE and ME-OC models have 213 DOF (225 time points - 12 regressors), represented by the blue horizontal line. Each point represents a scan modeled with ME-ICA (60 total). The black horizontal lines represent the lower quartile, median, and upper quartiles of the ME-ICA DOF.
